## Appendix S1 for "Rapid and specific degradation of endogenous proteins in mouse models using auxin-inducible degrons"

**Table S1A: Oligonucleotide details**

**Single Stranded Repair templates used for Easi-CRISPR. Exogenous sequences (linker:AID:Clover) are highlighted in green.**

***NCAPH* gRNA target sequence:** CCGCTGCAGACGTCTCAAAG

***NCAPH*-AID-mClover Repair Template ssODN**

TGTTCTCAAGTCTATCAAAGCCTGTGTTTCTTTCTTCTCTTCCCTCAGAATCTGAAGCTAGAAGGCACAGAGGATCTTTCTGATGTTCTGGTGATGCAAGGGGACGGGGGAGCAGGAGGCGTCGACGGTATCGATCCTAAAGATCCAGCCAAACCTCCGGCCAAGGCACAAGTTGTGGGATGGCCACCGGTGAGATCATACCGGAAGAACGTGATGGTTTCCTGCCAAAAATCAAGCGGTGGCCCGGAGGCGGCGGCGTTCGTGAAGAAGCTTGATGTGAGCAAGGGCGAGGAGCTGTTCACCGGGGTGGTGCCCATCCTGGTCGAGCTGGACGGCGACGTAAACGGCCACAAGTTCAGCGTCCGCGGCGAGGGCGAGGGCGATGCCACCAACGGCAAGCTGACCCTGAAGTTCATCTGCACCACCGGCAAGCTGCCCGTGCCCTGGCCCACCCTCGTGACCACCTTCGGCTACGGCGTGGCCTGCTTCAGCCGCTACCCCGACCACATGAAGCAGCACGACTTCTTCAAGTCCGCCATGCCCGAAGGCTACGTCCAGGAGCGCACCATCTCTTTCAAGGACGACGGTACCTACAAGACCCGCGCCGAGGTGAAGTTCGAGGGCGACACCCTGGTGAACCGCATCGAGCTGAAGGGCATCGACTTCAAGGAGGACGGCAACATCCTGGGGCACAAGCTGGAGTACAACTTCAACAGCCACAACGTCTATATCACGGCCGACAAGCAGAAGAACGGCATCAAGGCTAACTTCAAGATCCGCCACAACGTTGAGGACGGCAGCGTGCAGCTCGCCGACCACTACCAGCAGAACACCCCCATCGGCGACGGCCCCGTGCTGCTGCCCGACAACCACTACCTGAGCCATCAGTCCGCCCTGAGCAAAGACCCCAACGAGAAGCGCGATCACATGGTCCTGCTGGAGTTCGTGACCGCCGCCGGGATTACACATGGCATGGACGAGCTGTACAAGTGAGTACACTTTGAGACGTCTGCAGCGGGATGCCCATGGCTTCCTGGTGGCCGTGGAGAGGTGCTGGGCCTGTAGCGTGGCTATAGCCAACCACCACGGTGCTGG

***NCAPH2* gRNA target sequence:** ATCCATGGCCCAGCCTTGAG

***NCAPH2*-AID-mClover Repair Template ssODN**

CCAGGACTGGAGGCAGCTGTGGACACAATGTCTCTGAGACTGCTCACACACCAGCGAGCCCACACCCGCTTCCAGACCTATGCTGCACCATCCATGGCCCAGCCTGGGGGAGCAGGAGGCGTCGACGGTATCGATCCTAAAGATCCAGCCAAACCTCCGGCCAAGGCACAAGTTGTGGGATGGCCACCGGTGAGATCATACCGGAAGAACGTGATGGTTTCCTGCCAAAAATCAAGCGGTGGCCCGGAGGCGGCGGCGTTCGTGAAGAAGCTTGATGTGAGCAAGGGCGAGGAGCTGTTCACCGGGGTGGTGCCCATCCTGGTCGAGCTGGACGGCGACGTAAACGGCCACAAGTTCAGCGTCCGCGGCGAGGGCGAGGGCGATGCCACCAACGGCAAGCTGACCCTGAAGTTCATCTGCACCACCGGCAAGCTGCCCGTGCCCTGGCCCACCCTCGTGACCACCTTCGGCTACGGCGTGGCCTGCTTCAGCCGCTACCCCGACCACATGAAGCAGCACGACTTCTTCAAGTCCGCCATGCCCGAAGGCTACGTCCAGGAGCGCACCATCTCTTTCAAGGACGACGGTACCTACAAGACCCGCGCCGAGGTGAAGTTCGAGGGCGACACCCTGGTGAACCGCATCGAGCTGAAGGGCATCGACTTCAAGGAGGACGGCAACATCCTGGGGCACAAGCTGGAGTACAACTTCAACAGCCACAACGTCTATATCACGGCCGACAAGCAGAAGAACGGCATCAAGGCTAACTTCAAGATCCGCCACAACGTTGAGGACGGCAGCGTGCAGCTCGCCGACCACTACCAGCAGAACACCCCCATCGGCGACGGCCCCGTGCTGCTGCCCGACAACCACTACCTGAGCCATCAGTCCGCCCTGAGCAAAGACCCCAACGAGAAGCGCGATCACATGGTCCTGCTGGAGTTCGTGACCGCCGCCGGGATTACACATGGCATGGACGAGCTGTACAAGTAAGTGGACAGCACTGAGGCAGGGGTGGAAAGTAGTATATACCTGGAGGTCTTTGCCCCTAATGTGCTATGGGGCCATTCACTCCAGTGCTGCCTCCTGGCTGGCCTA

*Rosa26* sgRNA target sequence: GACTCCAGTCTTTCTAGAAGA

Sequence details for the donor template used to generate the LSL-Tir1-9myc insertion at Rosa26.

```
LOCUS       Exported               13430 bp DNA     linear   UNA 31-JUL-2019
DEFINITION  .
ACCESSION   .
VERSION     .
KEYWORDS    .
SOURCE      natural DNA sequence
  ORGANISM  unspecified
REFERENCE   1  (bases 1 to 13430)
  AUTHORS   .
  TITLE     Direct Submission
  JOURNAL   Exported Jan 10, 2022 from SnapGene Viewer 5.3.2
            https://www.snapgene.com
FEATURES             Location/Qualifiers
     source          1..13430
                     /organism="unspecified"
                     /mol_type="genomic DNA"
     misc_feature    271..1106
                     /locus_tag="R26 5arm"
                     /label=R26 5arm
                     /note="color: #00ffff"
     misc_feature    1096..1106
                     /locus_tag="5arm r"
                     /label=5arm r
                     /note="color: #00ffff"
     misc_feature    2019..3646
                     /locus_tag="CAG Promoter"
                     /label=CAG Promoter
                     /note="color: #ff001e"
     misc_feature    2019..2306
                     /locus_tag="CAG Enhancer"
                     /label=CAG Enhancer
                     /note="color: #00ffff"
     misc_feature    3652..3685
                     /locus_tag="LoxP"
                     /label=LoxP
                     /note="color: #00ffff"
     misc_feature    3686..3706
                     /locus_tag="SeqF"
                     /label=SeqF
                     /note="color: #fffd0f"
     misc_feature    3686..3693
                     /locus_tag="Loxp"
                     /label=Loxp
                     /note="color: #00ffff"
     misc_feature    3759..3766
                     /locus_tag="AsiSI"
                     /label=AsiSI
                     /note="color: #00ffff"
     misc_feature    3767..3772
                     /locus_tag="MluI"
                     /label=MluI
                     /note="color: #68ff25"
     misc_feature    3773..5629
                     /locus_tag="osTIR1"
                     /label=osTIR1
                     /note="color: #00ffff"
     misc_feature    5645..5650
                     /locus_tag="MluI(1)"
                     /label=MluI
                     /label=MluI(1)
                     /note="color: #68ff25"
     misc_feature    6200..7010
                     /locus_tag="R26 3arm"
                     /label=R26 3arm
                     /note="color: #00ffff"
     misc_feature    7011..7047
                     /locus_tag="3arm"
                     /label=3arm
                     /note="color: #00ffff"
ORIGIN
        1 caccgcatta ccctgttatc cctagcggca ggccctccga gcgtggtgga gccgttctgt
       61 gagacagccg ggtacgagtc gtgacgctgg aaggggcaag cgggtggtgg gcaggaatgc
      121 ggtccgccct gcagcaaccg gagggggagg gagaagggag cggaaaagtc tccaccggac
      181 gcggccatgg ctcggggggg ggggggcagc ggaggagcgc ttccggccga cgtctcgtcg
      241 ctgattggct tcttttcctc ccgccgtgtg tgaaaacaca aatggcgtgt tttggttggc
      301 gtaaggcgcc tgtcagttaa cggcagccgg agtgcgcagc cgccggcagc ctcgctctgc
      361 ccactgggtg gggcgggagg taggtggggt gaggcgagct ggacgtgcgg gcgcggtcgg
      421 cctctggcgg ggcgggggag gggagggagg gtcagcgaaa gtagctcgcg cgcgagcggc
      481 cgcccaccct ccccttcctc tgggggagtc gttttacccg ccgccggccg ggcctcgtcg
      541 tctgattggc tctcggggcc cagaaaactg gcccttgcca ttggctcgtg ttcgtgcaag
      601 ttgagtccat ccgccggcca gcgggggcgg cgaggaggcg ctcccaggtt ccggccctcc
      661 cctcggcccc gcgccgcaga gtctggccgc gcgcccctgc gcaacgtggc aggaagcgcg
      721 cgctgggggc ggggacgggc agtagggctg agcggctgcg gggcgggtgc aagcacgttt
      781 ccgacttgag ttgcctcaag aggggcgtgc tgagccagac ctccatcgcg cactccgggg
      841 agtggaggga aggagcgagg gctcagttgg gctgttttgg aggcaggaag cacttgctct
      901 cccaaagtcg ctctgagttg ttatcagtaa gggagctgca gtggagtagg cggggagaag
      961 gccgcaccct tctccggagg ggggagggga gtgttgcaat acctttctgg gagttctctg
     1021 ctgcctcctg gcttctgagg accgccctgg gcctgggaga atcccttccc cctcttccct
     1081 cgtgatctgc aactccagtc tttctaggcg cgcccgggct gcagatctgt agggcgcagt
     1141 agtccagggt ttccttgatg atgtcatact tatcctgtcc cttttttttc cacagctcgc
     1201 ggttgaggac aaactcttcg cggtctttcc agtggggatc gacggtatcg ataagctggc
     1261 cgctctagtg gccgtacggg cccacctgcc gggccactta attaaattta aatcacaatt
     1321 ccagctgagc gccggtcgct accattacca gttggtctgg tgtcaaaaat aataataacc
     1381 gggcaggggg gatctgcatg gatctttgtg aaggaacctt acttctgtgg tgtgacataa
     1441 ttggacaaac actccgaggc ggatcacaag catacctaca gagatttaaa gctctaaggt
     1501 aaatataaaa tttttaagtg tataatgtgt taaactactg attctaattg tttgtgtatt
     1561 ttagattcca acctatggaa ctgatgaatg ggagcagtgg tggaatgcca gatccagaca
     1621 tgataagata cattgatgag tttggacaaa ccacaactag aatgcagtga aaaaaatgct
     1681 ttatttgtga aatttgtgat gctattgctt tatttgtaac cattataagc tgcaataaac
     1741 aagttaacaa caacaattgc attcatttta tgtttcaggt tcagggggag gtgtgggagg
     1801 ttttttaaag caagtaaaac ctctacaaat gtggtatggc tgattatgat ctgcggccgt
     1861 gctagcgctt aagcttgaag ttcctattcc gaagttccta ttctctagaa agtataggaa
     1921 cttcggcgcg ccggatctga cattgattat tgactagtta ttaatagtaa tcaattacgg
     1981 ggtcattagt tcatagccca tatatggagt tccgcgttac ataacttacg gtaaatggcc
     2041 cgcctggctg accgcccaac gacccccgcc cattgacgtc aataatgacg tatgttccca
     2101 tagtaacgcc aatagggact ttccattgac gtcaatgggt ggagtattta cggtaaactg
     2161 cccacttggc agtacatcaa gtgtatcata tgccaagtac gccccctatt gacgtcaatg
     2221 acggtaaatg gcccgcctgg cattatgccc agtacatgac cttatgggac tttcctactt
     2281 ggcagtacat ctacgtatta gtcatcgcta ttaccatggg tcgaggtgag ccccacgttc
     2341 tgcttcactc tccccatctc ccccccctcc ccacccccaa ttttgtattt atttattttt
     2401 taattatttt gtgcagcgat gggggcgggg gggggggggg cgcgcgccag gcggggcggg
     2461 gcggggcgag gggcggggcg gggcgaggcg gagaggtgcg gcggcagcca atcagagcgg
     2521 cgcgctccga aagtttcctt ttatggcgag gcggcggcgg cggcggccct ataaaaagcg
     2581 aagcgcgcgg cgggcgggag tcgctgcgtt gccttcgccc cgtgccccgc tccgcgccgc
     2641 ctcgcgccgc ccgccccggc tctgactgac cgcgttactc ccacaggtga gcgggcggga
     2701 cggcccttct cctccgggct gtaattagcg cttggtttaa tgacggctcg tttcttttct
     2761 gtggctgcgt gaaagcctta aagggctccg ggagggccct ttgtgcgggg gggagcggct
     2821 cggggggtgc gtgcgtgtgt gtgtgcgtgg ggagcgccgc gtgcggcccg cgctgcccgg
     2881 cggctgtgag cgctgcgggc gcggcgcggg gctttgtgcg ctccgcgtgt gcgcgagggg
     2941 agcgcggccg ggggcggtgc cccgcggtgc gggggggctg cgaggggaac aaaggctgcg
     3001 tgcggggtgt gtgcgtgggg gggtgagcag ggggtgtggg cgcggcggtc gggctgtaac
     3061 ccccccctgc acccccctcc ccgagttgct gagcacggcc cggcttcggg tgcggggctc
     3121 cgtgcggggc gtggcgcggg gctcgccgtg ccgggcgggg ggtggcggca ggtgggggtg
     3181 ccgggcgggg cggggccgcc tcgggccggg gagggctcgg gggaggggcg cggcggcccc
     3241 ggagcgccgg cggctgtcga ggcgcggcga gccgcagcca ttgcctttta tggtaatcgt
     3301 gcgagagggc gcagggactt cctttgtccc aaatctggcg gagccgaaat ctgggaggcg
     3361 ccgccgcacc ccctctagcg ggcgcgggcg aagcggtgcg gcgccggcag gaaggaaatg
     3421 ggcggggagg gccttcgtgc gtcgccgcgc cgccgtcccc ttctccatct ccagcctcgg
     3481 ggctgccgca gggggacggc tgccttcggg ggggacgggg cagggcgggg ttcggcttct
     3541 ggcgtgtgac cggcggctct agagcctctg ctaaccatgt tcatgccttc ttctttttcc
     3601 tacagatcct taattaaccg tcttaagcct gtaacaaccg gtacagttcg aataacttcg
     3661 tatagcatac attatacgaa gttatgaagt tataccgggc caccatagtc gcgagtagct
     3721 tgggccagct aggccttgac caaagttcct ctggaattgc gatcgcacgc gtatgacgta
     3781 cttcccggag gaggtggtgg agcacatctt cagcttcctg ccggcgcagc gcgaccgcaa
     3841 cacggtctcg ctcgtctgca aggtgtggta cgagatcgag aggctgagcc gccgcggcgt
     3901 cttcgtgggc aactgctacg ccgtgcgcgc cggccgcgtc gccgcgcggt tccccaacgt
     3961 gcgggcgctc acggtgaagg ggaagcccca cttcgccgac ttcaacctcg tgccccccga
     4021 ctggggcggc tacgcggggc cgtggatcga ggcggccgcg aggggatgcc acggcctgga
     4081 ggagctcagg atgaagcgga tggtggtgtc cgacgagagc ctcgagctgc tggctcgctc
     4141 gttcccgcgg ttcagggctc ttgttcttat cagctgcgag gggttcagca ctgacgggct
     4201 agccgccgtc gcgagccatt gcaagcttct gagggagttg gatttgcagg aaaatgaagt
     4261 ggaggatcga gggcctaggt ggctttcctg cttccctgat tcctgcacat cacttgtctc
     4321 attgaatttt gcctgcatca aaggggaggt taatgctggt tcactggaga gacttgttag
     4381 caggtcccca aacctgcgga gtttgaggct gaatcgatct gtatcggtag atacacttgc
     4441 aaagatacta ctgcgtaccc ctaacttgga ggatttgggg acagggaatt tgacagatga
     4501 cttccaaact gagtcctact ttaagcttac cagtgctctg gagaaatgca agatgttgag
     4561 gagtttgtct ggattctggg atgcttctcc tgtttgcctg tcatttatct accccctgtg
     4621 tgctcaactg acaggattga acttgagcta tgcacccaca cttgatgctt ctgaccttac
     4681 aaaaatgatt agccgctgtg tgaagctcca acgcctttgg gtactggatt gtatctcgga
     4741 caaaggcttg caagtggtgg cctccagttg caaagacttg caagaactca gggtatttcc
     4801 atcagatttc tacgtagctg gttattctgc agtgacagag gagggacttg ttgcagtatc
     4861 cttgggctgt ccaaaactga actcactact gtacttctgt caccaaatga ctaatgctgc
     4921 actagttact gtcgccaaga actgtccaaa tttcacacga ttcagacttt gtattcttga
     4981 gccagggaag cctgatgttg tgacaagcca accattagat gaaggctttg gagctattgt
     5041 tcgtgagtgc aagggattac aacgtttgtc aatatctggt cttctcacag acaaagtttt
     5101 catgtatatt gggaaatatg caaaacaact tgagatgctt tctatagcat ttgctggtga
     5161 cagtgataag ggtatgatgc atgttatgaa tggatgcaag aatttaagga aactggagat
     5221 aagagatagc ccgtttggtg atgctgcact cttggggaat tttgctaggt acgagacaat
     5281 gcgatccctt tggatgtcat cttgcaatgt cacgttaaag gggtgccaag tccttgcgtc
     5341 aaagatgccg atgctcaatg ttgaggtcat aaatgagcgg gatggtagca atgaaatgga
     5401 ggaaaaccat ggagatctgc ctaaagtgga gaaattatat gtgtaccgca caactgctgg
     5461 ggcgagggat gatgcaccaa attttgttaa aatcctagtc gactccggtt ctgctgctag
     5521 tggtgaacaa aagttgattt ctgaagaaga tttgaacggt gaacaaaagc taatctccga
     5581 ggaagacttg aacggtgaac aaaaattaat ctcagaagaa gacttgaacg gatccactag
     5641 ctaaacgcgt aaatgattgc agatccacta gttctagagc tcgctgatca gcctcgactg
     5701 tgccttctag ttgccagcca tctgttgttt gcccctcccc cgtgccttcc ttgaccctgg
     5761 aaggtgccac tcccactgtc ctttcctaat aaaatgagga aattgcatcg cattgtctga
     5821 gtaggtgtca ttctattctg gggggtgggg tggggcagga cagcaagggg gaggattggg
     5881 aagacaatag caggcatgct ggggatgcgg tgggctctat ggcttctgag gcggaaagaa
     5941 ccagctgggg ctcgagcctc tagaactata gtgagtcgta ttacgtagat ccagacatga
     6001 taagatacat tgatgagttt ggacaaacca caactagaat gcagtgaaaa aaatgcttta
     6061 tttgtgaaat ttgtgatgct attgctttat ttgtaaccat tataagctgc aataaacaag
     6121 ttaacaacaa caattgcatt cattttatgt ttcaggttca gggggaggtg tgggaggttt
     6181 tttaattcgc ggccctagaa gatgggcggg agtcttctgg gcaggcttaa aggctaacct
     6241 ggtgtgtggg cgttgtcctg caggggaatt gaacaggtgt aaaattggag ggacaagact
     6301 tcccacagat tttcggtttt gtcgggaagt tttttaatag gggcaaataa ggaaaatggg
     6361 aggataggta gtcatctggg gttttatgca gcaaaactac aggttattat tgcttgtgat
     6421 ccgcctcgga gtattttcca tcgaggtaga ttaaagacat gctcacccga gttttatact
     6481 ctcctgcttg agatccttac tacagtatga aattacagtg tcgcgagtta gactatgtaa
     6541 gcagaatttt aatcattttt aaagagccca gtacttcata tccatttctc ccgctccttc
     6601 tgcagcctta tcaaaaggta ttttagaaca ctcattttag ccccattttc atttattata
     6661 ctggcttatc caacccctag acagagcatt ggcattttcc ctttcctgat cttagaagtc
     6721 tgatgactca tgaaaccaga cagattagtt acatacacca caaatcgagg ctgtagctgg
     6781 ggcctcaaca ctgcagttct tttataactc cttagtacac tttttgttga tcctttgcct
     6841 tgatccttaa ttttcagtgt ctatcacctc tcccgtcagg tggtgttcca catttgggcc
     6901 tattctcagt ccagggagtt ttacaacaat agatgtattg agaatccaac ctaaagctta
     6961 actttccact cccatgaatg cctctctcct ttttctccat ttataaactg agctattaac
     7021 cattaatggt ttccaggtgg atgtctcctc ccccaatatt acctgatgta tcttacatat
     7081 tgccaggctg atattttaag acattaaaag gtatatttca ttattgagcc acatggtatt
     7141 gattactgct tactaaaatt ttgtcattgt acacatctgt aaaaggtggt tccttttgga
     7201 atgcaaagtt caggtgtttg ttgtctttcc tgacctaagg tcttgtgagc ttgtattttt
     7261 tctatttaag cagtgctttc tcttggactg gcttgactca tggcattcta cacgttattg
     7321 ctggtctaaa tgtgattttg ccaagcttct tcaggaccta taattttgct tgacttgtag
     7381 ccaaacacaa gtaaaatgat taagcaacaa atgtatttgt gaagcttggt ttttaggttg
     7441 ttgtgttgtg tgtgcttgtg ctctataata atactatcca ggggctggag aggtggctcg
     7501 gagttcaaga gcacagactg ctcttccaga agtcctgagt tcaattccca gcaaccacat
     7561 ggtggctcac aaccatctgt aatgggatct gatgccctct tctggtgtgt ctgaagacca
     7621 caagtgtatt cacattaaat aaataaatcc tccttcttct tctttttttt ttttttaaag
     7681 agaatactgt ctccagtaga atttactgaa gtaatgaaat actttgtgtt tgttccaata
     7741 tggtagccaa taatcaaatt actctttaag cactggaaat gttaccaagg aactaatttt
     7801 tatttgaagt gtaactgtgg acagaggagc cataactgca gacttgtggg atacagaaga
     7861 ccaatgcaga ctttaatgtc ttttctctta cactaagcaa taaagaaata aaaattgaac
     7921 ttctagtatc ctatttgttt aaactgctag ctttacttaa cttttgtgct tcatctatac
     7981 aaagctgaaa gctaagtctg cagccattac taaacatgaa agcaagtaat gataattttg
     8041 gatttcaaaa atgtagggcc agagtttagc cagccagtgg tggtgcttgc ctttatgcct
     8101 ttaatcccag cactctggag gcagagacag gcagatctct gagtttgagc ccagcctggt
     8161 ctacacatca agttctatct aggatagcca ggaatacaca cagaaaccct gttggggagg
     8221 ggggctctga gatttcataa aattataatt gaagcattcc ctaatgagcc actatggatg
     8281 tggctaaatc cgtctacctt tctgatgaga tttgggtatt attttttctg tctctgctgt
     8341 tggttgggtc ttttgacact gtgggctttc tttaaagcct ccttcctgcc atgtggtctc
     8401 ttgtttgcta ctaacttccc atggcttaaa tggcatggct ttttgccttc taagggcagc
     8461 tgctgagatt tgcagcctga tttccagggt ggggttggga aatctttcaa acactaaaat
     8521 tgtcctttaa tttttttttt aaaaaatggg ttatataata aacctcataa aatagttatg
     8581 aggagtgagg tggactaata ttaaatgagt ccctccccta taaaagagct attaaggctt
     8641 tttgtcttat acttaacttt ttttttaaat gtggtatctt tagaaccaag ggtcttagag
     8701 ttttagtata cagaaactgt tgcatcgctt aatcagattt tctagtttca aatccagaga
     8761 atccaaattc ttcacagcca aagtcaaatt aagaatttct gacttttaat gttaatttgc
     8821 ttactgtgaa tataaaaatg atagcttttc ctgaggcagg gtctcactat gtatctctgc
     8881 ctgatctgca acaagatatg tagactaaag ttctgcctgc ttttgtctcc tgaatactaa
     8941 ggttaaaatg tagtaatact tttggaactt gcaggtcaga ttcttttata ggggacacac
     9001 taagggagct tgggtgatag ttggtaaaat gtgtttcaag tgatgaaaac ttgaattatt
     9061 atcaccgcaa cctacttttt aaaaaaaaaa gccaggcctg ttagagcatg cttaagggat
     9121 ccctaggact tgctgagcac acaagagtag ttacttggca ggctcctggt gagagcatat
     9181 ttcaaaaaac aaggcagaca accaagaaac tacagttaag gttacctgtc tttaaaccat
     9241 ctgcatatac acagggatat taaaatattc caaataatat ttcattcaag ttttccccca
     9301 tcaaattggg acatggattt ctccggtgaa taggcagagt tggaaactaa acaaatgttg
     9361 gttttgtgat ttgtgaaatt gttttcaagt gatagttaaa gcccatgaga tacagaacaa
     9421 agctgctatt tcgaggtctc ttggtttata ctcagaagca cttctttggg tttccctgca
     9481 ctatcctgat catgtgctag gcctacctta ggctgattgt tgttcaaata aacttaagtt
     9541 tcctgtcagg tgatgtcata tgatttcata tatcaaggca aaacatgtta tatatgttaa
     9601 acatttgtac ttaatgtgaa agttaggtct ttgtgggttt gatttttaat tttcaaaacc
     9661 tgagctaaat aagtcatttt tacatgtctt acatttggtg gaattgtata attgtggttt
     9721 gcaggcaaga ctctctgacc tagtaaccct acctatagag cactttgctg ggtcacaagt
     9781 ctaggagtca agcatttcac cttgaagttg agacgttttg ttagtgtata ctagtttata
     9841 tgttggagga catgtttatc cagaagatat tcaggactat ttttgactgg gctaaggaat
     9901 tgattctgat tagcactgtt agtgagcatt gagtggcctt taggcttgaa ttggagtcac
     9961 ttgtatatct caaataatgc tggccttttt taaaaagccc ttgttcttta tcaccctgtt
    10021 ttctacataa tttttgttca aagaaatact tgtttggatc tccttttgac aacaatagca
    10081 tgttttcaag ccatattttt tttccttttt tttttttttt ttggtttttc gagacagggt
    10141 ttctctgtat agccctggct gtcctggaac tcactttgta gaccaggctg gcctcgaact
    10201 cagaaatccg cctgcctctg cctcctgagt gccgggatta aaggcgtgca ccaccacgcc
    10261 tggctaagtt ggatattttg ttatataact ataaccaata ctaactccac tgggtggatt
    10321 tttaattcag tcagtagtct taagtggtct ttattggccc ttcattaaaa tctactgttc
    10381 actctaacag aggctgttgg tactagtggc acttaagcaa cttcctacgg atatactagc
    10441 agattaaggg tcagggatag aaactagtct agcgttttgt atacctacca gctttatact
    10501 accttgttct gatagaaata tttcaggaca tctagcttat cgataccgtc gacggtatcg
    10561 ataagcttga tccagctttt gttcccttta gtgagggtta attgcgcgct tggcgtaatc
    10621 atggtcatag ctgtttcctg tgtgaaattg ttatccgctc acaattccac acaacatacg
    10681 agccggaagc ataaagtgta aagcctgggg tgcctaatga gtgagctaac tcacattaat
    10741 tgcgttgcgc tcactgcccg ctttccagtc gggaaacctg tcgtgccagc tgcattaatg
    10801 aatcggccaa cgcgcgggga gaggcggttt gcgtattggg cgctcttccg cttcctcgct
    10861 cactgactcg ctgcgctcgg tcgttcggct gcggcgagcg gtatcagctc actcaaaggc
    10921 ggtaatacgg ttatccacag aatcagggga taacgcagga aagaacatgt gagcaaaagg
    10981 ccagcaaaag gccaggaacc gtaaaaaggc cgcgttgctg gcgtttttcc ataggctccg
    11041 cccccctgac gagcatcaca aaaatcgacg ctcaagtcag aggtggcgaa acccgacagg
    11101 actataaaga taccaggcgt ttccccctgg aagctccctc gtgcgctctc ctgttccgac
    11161 cctgccgctt accggatacc tgtccgcctt tctcccttcg ggaagcgtgg cgctttctca
    11221 tagctcacgc tgtaggtatc tcagttcggt gtaggtcgtt cgctccaagc tgggctgtgt
    11281 gcacgaaccc cccgttcagc ccgaccgctg cgccttatcc ggtaactatc gtcttgagtc
    11341 caacccggta agacacgact tatcgccact ggcagcagcc actggtaaca ggattagcag
    11401 agcgaggtat gtaggcggtg ctacagagtt cttgaagtgg tggcctaact acggctacac
    11461 tagaaggaca gtatttggta tctgcgctct gctgaagcca gttaccttcg gaaaaagagt
    11521 tggtagctct tgatccggca aacaaaccac cgctggtagc ggtggttttt ttgtttgcaa
    11581 gcagcagatt acgcgcagaa aaaaaggatc tcaagaagat cctttgatct tttctacggg
    11641 gtctgacgct cagtggaacg aaaactcacg ttaagggatt ttggtcatga gattatcaaa
    11701 aaggatcttc acctagatcc ttttaaatta aaaatgaagt tttaaatcaa tctaaagtat
    11761 atatgagtaa acttggtctg acagttacca atgcttaatc agtgaggcac ctatctcagc
    11821 gatctgtcta tttcgttcat ccatagttgc ctgactcccc gtcgtgtaga taactacgat
    11881 acgggagggc ttaccatctg gccccagtgc tgcaatgata ccgcgagacc cacgctcacc
    11941 ggctccagat ttatcagcaa taaaccagcc agccggaagg gccgagcgca gaagtggtcc
    12001 tgcaacttta tccgcctcca tccagtctat taattgttgc cgggaagcta gagtaagtag
    12061 ttcgccagtt aatagtttgc gcaacgttgt tgccattgct acaggcatcg tggtgtcacg
    12121 ctcgtcgttt ggtatggctt cattcagctc cggttcccaa cgatcaaggc gagttacatg
    12181 atcccccatg ttgtgcaaaa aagcggttag ctccttcggt cctccgatcg ttgtcagaag
    12241 taagttggcc gcagtgttat cactcatggt tatggcagca ctgcataatt ctcttactgt
    12301 catgccatcc gtaagatgct tttctgtgac tggtgagtac tcaaccaagt cattctgaga
    12361 atagtgtatg cggcgaccga gttgctcttg cccggcgtca atacgggata ataccgcgcc
    12421 acatagcaga actttaaaag tgctcatcat tggaaaacgt tcttcggggc gaaaactctc
    12481 aaggatctta ccgctgttga gatccagttc gatgtaaccc actcgtgcac ccaactgatc
    12541 ttcagcatct tttactttca ccagcgtttc tgggtgagca aaaacaggaa ggcaaaatgc
    12601 cgcaaaaaag ggaataaggg cgacacggaa atgttgaata ctcatactct tcctttttca
    12661 atattattga agcatttatc agggttattg tctcatgagc ggatacatat ttgaatgtat
    12721 ttagaaaaat aaacaaatag gggttccgcg cacatttccc cgaaaagtgc cacctaaatt
    12781 gtaagcgtta atattttgtt aaaattcgcg ttaaattttt gttaaatcag ctcatttttt
    12841 aaccaatagg ccgaaatcgg caaaatccct tataaatcaa aagaatagac cgagataggg
    12901 ttgagtgttg ttccagtttg gaacaagagt ccactattaa agaacgtgga ctccaacgtc
    12961 aaagggcgaa aaaccgtcta tcagggcgat ggcccactac gtgaaccatc accctaatca
    13021 agttttttgg ggtcgaggtg ccgtaaagca ctaaatcgga accctaaagg gagcccccga
    13081 tttagagctt gacggggaaa gccggcgaac gtggcgagaa aggaagggaa gaaagcgaaa
    13141 ggagcgggcg ctagggcgct ggcaagtgta gcggtcacgc tgcgcgtaac caccacaccc
    13201 gccgcgctta atgcgccgct acagggcgcg tcccattcgc cattcaggct gcgcaactgt
    13261 tgggaagggc gatcggtgcg ggcctcttcg ctattacgcc agctggcgaa agggggatgt
    13321 gctgcaaggc gattaagttg ggtaacgcca gggttttccc agtcacgacg ttgtaaaacg
    13381 acggccagtg agcgcgcgta atacgactca ctatagggcg aattggagct
//
```
